# The thermodynamics of reproduction constrain species ranking dynamics and diversity

**DOI:** 10.1101/2022.08.06.503003

**Authors:** Tom Froese, Rainer Froese

## Abstract

Iñiguez, Pineda, Gershenson, and Barabási proposed that complex systems can be simplified and analyzed in terms of the dynamics of ordered lists. For social, economic, and infrastructural systems, they succeeded in modeling the dynamics of ranking with two mechanisms, namely displacement (i.e., shuffling of ranks among existing elements) and replacement of elements by new elements. A general pattern is that, for open systems with influx and outflux of elements, e.g., a list with a subset of elements, only the top of the list is stable; for closed systems, top and bottom are stable. This model was fitted to empirical data, resulting in a universal curve in which displacement and replacement parameters are inversely related, which implies that a single parameter could regulate both. Our aim is twofold: First, we demonstrate that the pattern generalizes to ranking of biological species, based on FishBase records of fish taxonomy and life history traits. Second, we propose a candidate for a unified mechanism, based on a recent model of the thermodynamics of self-replication.

## Main text

Iñiguez, Pineda, Gershenson, and Barabási ^1^ proposed that complex systems can be simplified and analyzed in terms of the dynamics of ordered lists. For social, economic, and infrastructural systems, they succeeded in modeling the dynamics of ranking with two mechanisms, namely displacement (i.e., shuffling of ranks among existing elements) and replacement of elements by new elements. A general pattern is that, for open systems with influx and outflux of elements, e.g., a list with a subset of elements, only the top of the list is stable; for closed systems, top and bottom are stable. This model was fitted to empirical data, resulting in a universal curve in which displacement and replacement parameters are inversely related, which implies that a single parameter could regulate both. Our aim is twofold: First, we demonstrate that the pattern generalizes to ranking of biological species, based on FishBase records of fish taxonomy and life history traits ^2^. Second, we propose a candidate for a unified mechanism, based on a recent model of the thermodynamics of self-replication ^3^.

In organisms that grow throughout their lives, such as invertebrates and fish, maximum body weight (*W*_*max*_) is a suitable variable for ranking; it is a proxy for many other variables of ecological and evolutionary importance ^4^, but which are more difficult to observe, such as generation time ^5^. We ranked 31,942 fish species with estimates of *W*_*max*_, representing over 90% of known fish species ^2^, which makes this list approximate a closed system. They exhibit *W*_*max*_ ranging over ten orders of magnitude, from 0.004 grams to 34 tons, with a median of 31.5 g, representing a typical fish of about 15 cm length. We calculated the proportional difference between ranks as follows: (*W*_*max*_ of rank[x] - *W*_*max*_ of rank[x+1]) / *W*_*max*_ of rank[x], giving a range from 0 (no difference) to 1 (a difference of *W*_*max*_).

We found that the proportional difference between ranks is dependent on rank, with the largest differences at the top and bottom of the list (Fig. 1). Compared to most species, these exceptionally large and small species have a *W*_*max*_ that is more distinct from that of adjacently ranked species, which entails that displacement from their ranks will be less frequent. This is consistent with Iñiguez et al.’s general pattern that lists of closed systems are equally stable at top and bottom ^1^. We could arbitrarily create a list of an open system with a subset of species, for example half of the top species ranked according to largest *W*_*max*_ (or, alternatively, ranked according to smallest *W*_*max*_ to be representative of shortest generation time), in which case only the top of the list would be stable.

**Figure 1.**
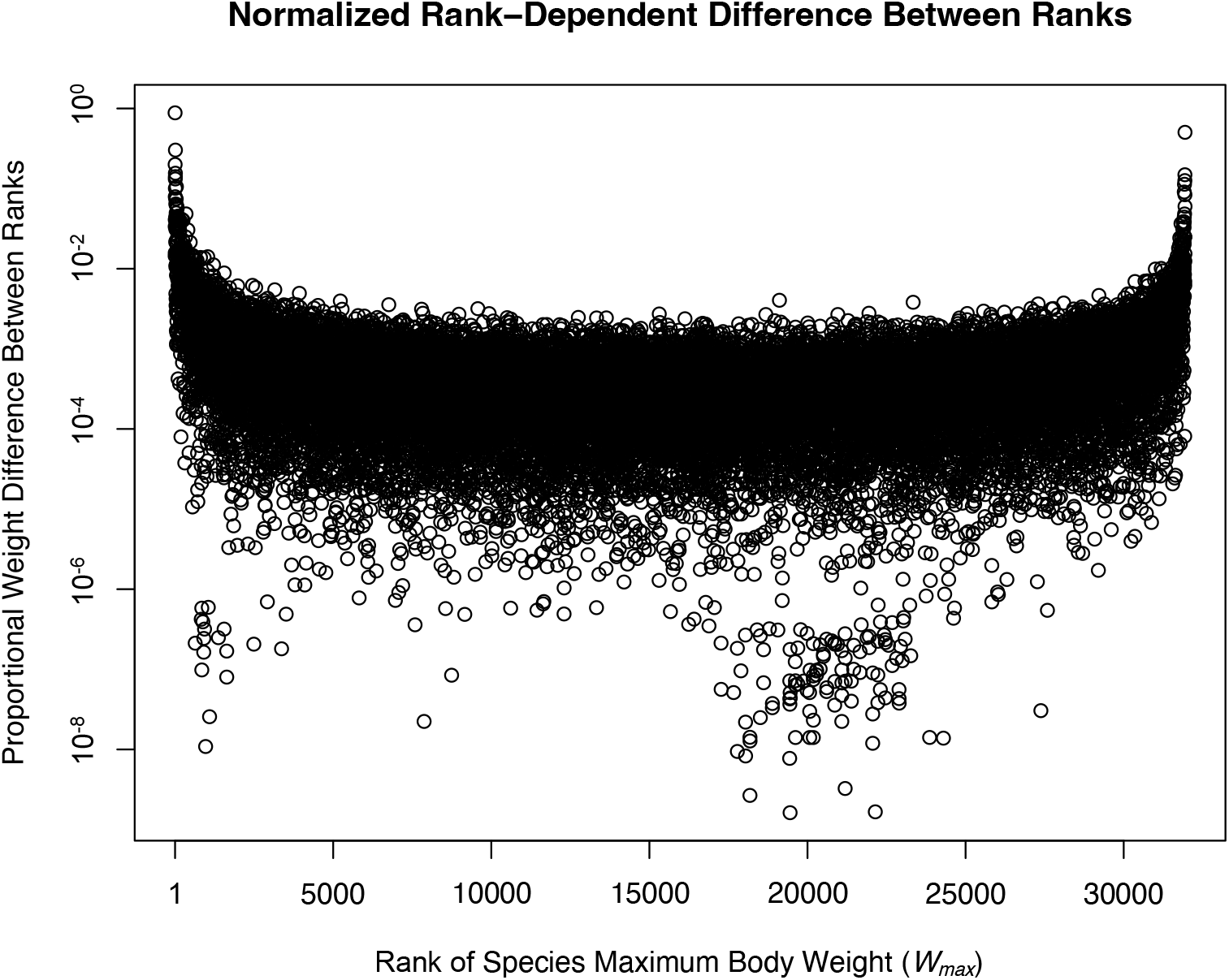
Rank difference (normalized range [0, 1]) between 31,942 fish species ranked by maximum body weight. For each fish species the proportional weight difference between two consecutive ranks was plotted, i.e., the difference in weight between one rank and the next lower rank divided by the weight of the higher rank. Top and bottom ranks are characterized by the largest weight differences between ranks, which entails that exceptionally large and small species are less likely to change ranks compared to most other species.

The finding that fish species with the largest *W*_*max*_ are associated with rank stability is consistent with known biology. In a submitted study ^6^, we show that fish species tend to be arranged along an evolutionary axis from large, little productive, and evolutionarily older species to small, highly productive, and evolutionarily younger species. Biologists explain this evolutionary pattern in terms of several domain-specific mechanisms: species with larger *W*_*max*_ speciate less rapidly because they tend to have longer generation times, and because they also tend to have smaller populations harboring less genetic diversity ^2^. And yet, given the fact that this biological example of rank stability is consistent with a general pattern of rank dynamics, there may be a domain-independent overarching principle at work as well.

We propose that thermodynamics can provide the theoretical framework from which to derive such a principle, specifically from the statistical physics of self-replication. A well-known insight in that field is that a replicator with net growth must have a positive lower bound on the total entropy production associated with its replication ^7^. If such a replicator is to replace another replicator under steady state conditions, the fitness difference of the former compared to the latter must be larger than a critical selection coefficient (*s*, range [0, 1], where 0 means that replacement is possible with minimal fitness difference). Recently, it has been shown that *s* is a function of the energetics of replication events ^3^, namely that its lower bound is given by the exponential of the negative of the total Gibbs free energy dissipated, i.e., *s* ≥*e*^*-ΔG*^. In other words, the more a replicator dissipates free energy during self-replication, the less fitness difference is required for it to replace another species.

We consider *W*_*max*_ as an indicator of maximum free energy dissipated in a reproduction event ^8^ as well as for all reproduction during the whole lifespan. Over evolutionary timescales, an approximately 1-to-1 ratio of body mass and self-replication dissipation of free energy (*W*_*max*_ ≈Δ*G*) makes sense: in a population at steady state, one female will on average be replaced by one new female ^9^. We therefore derive the following hypothesis: *the free energy dissipated during lifetime reproduction sets an upper bound on a species’ phylogenetic uniqueness*. Given that the critical selection coefficient *s* follows an exponential decay with increasing dissipation, *e*^*-ΔG*^, this upper bound is expected to apply to species with the smallest *W*_*max*_. Conversely, species with larger *W*_*max*_ tend to require less fitness difference to replace similar and newly diverging lineages, and hence can also occupy less diverse areas of phylogenetic space.

To test this hypothesis with the fish data, we analyzed species’ phylogenetic uniqueness using a standard scale of phylogenetic diversity (PD50) ^10^, given that high values of PD50 indicate the absence of phylogenetic diversity (such as whale shark *Rhincodon typus* being the only species in its Genus and Family, with PD50 = 1.5). As hypothesized, *W*_*max*_, taken as indicative of the lifetime reproductive dissipation of free energy, *ΔG*, is associated with an upper bound on phylogenetic uniqueness (Fig. 2). This values of PD50 quickly increase with *W*_*max*_ until some species regularly attain the highest level of uniqueness. *Dentatherina merceri* is the species with the smallest *W*_*max*_ (1.1 g) to achieve this uniqueness (PD50 = 1.5). Note there are two outliers with a PD50 >= 2, *Amia calva* and *Neoceratodus forsteri*, which are considered a relic and living fossil, respectively. Given that a maximum critical selection coefficient, *s* = 1, predicts the lowest upper bound on phylogenetic uniqueness (PD50 = 0.5), while conversely *s* = 0 predicts the highest upper bound (PD50 ≈ 2), and approximating *W*_*max*_ ≈ *ΔG*, we calculated this upper bound on PD50 as: (1 - *e*^*-Wmax*^) * (2 – 0.5) + 0.5. Most species’ PD50 falls under this upper bound; that some very small species remain above the bound on uniqueness could be an indication that there are closely related species waiting to be discovered.

**Figure 2:**
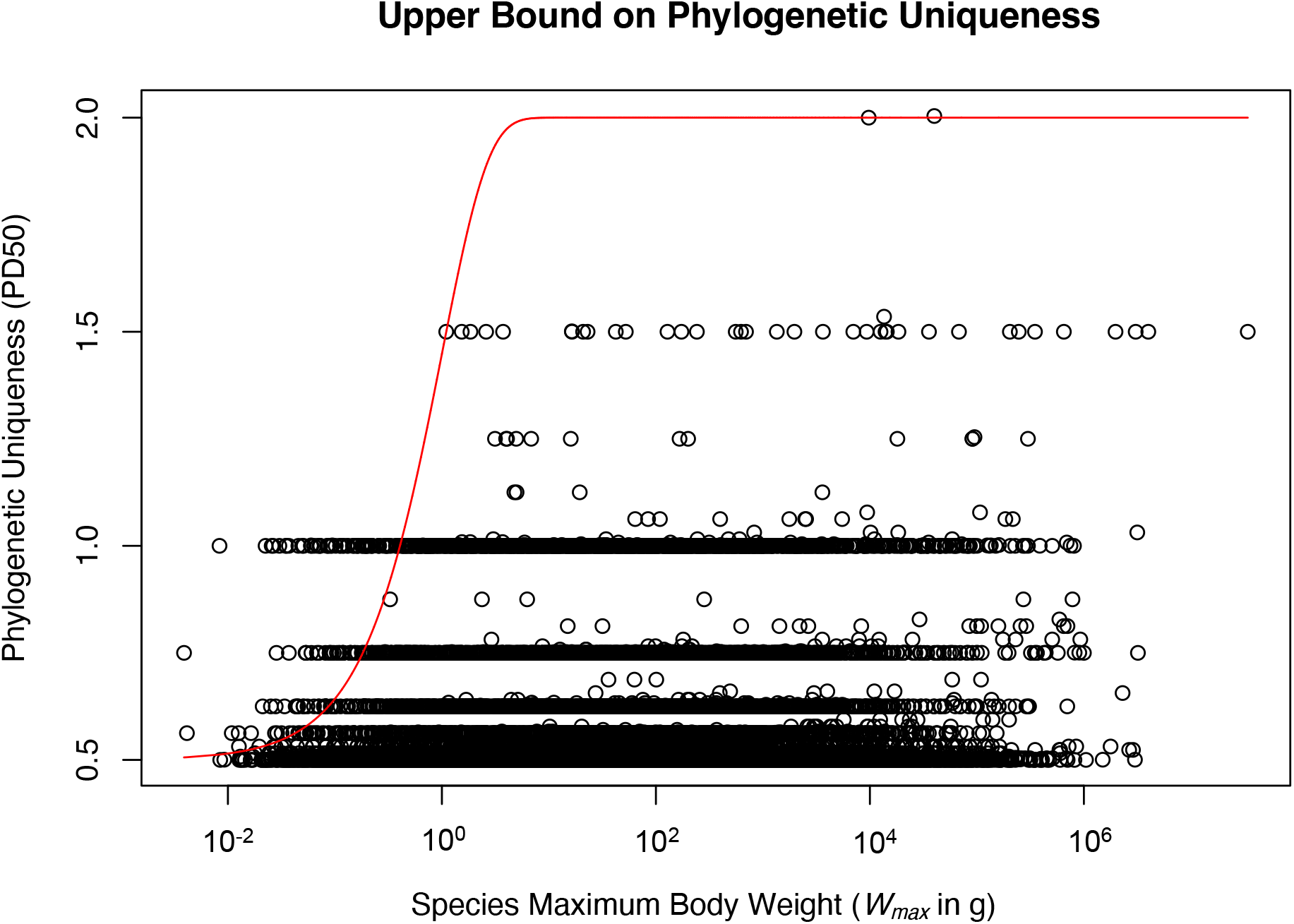
Phylogenetic uniqueness (using PD50 scale ^10^) over maximum body weight (*W*_*max*_) for 31,942 species of fishes. A PD50 of 0.5 indicates the highest phylogenetic diversity, with many closely related species in the same Genus, while a PD50 >= 1.5 indicates highest phylogenetic uniqueness with one or very few species in the same Order or Class. The red curve represents the upper bound on phylogenetic uniqueness (PD50) predicted from the critical selection coefficient (*s*) as a function of free energy dissipated (*e*^*-ΔG*^) approximating *ΔG* by *W*_*max*_.

The thermodynamic model also makes a prediction about the relationship between selection pressure and replacement dynamics ^3^: with increasing selection pressure, a replicator’s rate of replication becomes more important than *ΔG*, and replicators with faster rates will replace those with slower rates. This is consistent with the fact that fish populations respond to selection pressure from overfishing by evolving towards shorter generation time and smaller *W*_*max*_^11^. Species that naturally have the smallest *W*_*max*_ also compete by means of exceptionally short generation times, i.e., by maximizing the intrinsic rate of population growth rather than lifetime reproductive output. They achieve this by reproducing while they still have body characteristics of early juveniles or late larvae. Indeed, differences in rates of population growth link back to differences in *W*_*max*_: net growth sets a positive lower bound on total entropy produced (*ΔS*), and hence on *ΔG*, during reproduction ^7^. In other words, given a fixed or limited amount of free energy (*G*), the smallest species may achieve higher intrinsic rates of population growth by accommodating higher entropy production (*ΔS*) with lower energy investment (*ΔE*) in body growth (given that *G* = *E* – *TS*), resulting in underdeveloped *W*_*max*_.

Intriguingly, the thermodynamic model’s two distinct modes of replacement dynamics, namely dissipation-vs. rate-dependent, appear to fit the clustering of open systems on Iñiguez et al.’s universal curve: those systems with parameters that place them closer to what they call the “replacement regime” are characterized by higher energetic costs (e.g., cities, competitive sports, and academic rankings). Systems with parameters that place them closer to the other end tend to consist in elements that have lower energetic costs and faster turn-over, such as communicative acts in digital media (e.g., views, likes, comments) and print media (e.g., word choices in different languages). If this clustering can be substantiated, then the universal curve could be developed into a unified list of open systems ordered by the free energy dissipation required by elements to stay in top place.

## Author Contributions Statement

TF conceived of the idea and wrote the first draft of the manuscript. RF analyzed the data. TF and RF finalized the manuscript together.

## Acknowledgments

We thank Andrés G. Mejía Ramón for help in generating the figures. The manuscript benefited from discussions with Gerardo Iñiguez, Carlos Pineda, and Andrés G. Mejía Ramón and from feedback following its presentation at the GEOMAR Helmholtz Centre for Ocean Research.

